# Evaluation of the effect of SiO2 and Fe3O4 nanoparticles on *Pisum sativum* seeds in laboratory and field experiments

**DOI:** 10.1101/2020.08.31.275859

**Authors:** L.V. Galaktionova, A.M. Korotkova, N.I. Voskobulova, S.V. Lebedev, N.A. Terehova, I.A. Vershinina

## Abstract

The present study assessed the toxic effects and prospects of using nanoparticles of SiO2 and Fe3O4 by studying the influence of pre-sowing priming of *Pisum sativum L*. seeds with a suspension of nanoparticles of SiO_2_ and Fe_3_O_4_ in a concentration range of 10^−2^ to 10^−5^ mg/l. The results demonstrated the stimulating effect of the SiO_2_ suspension (10^−3^ mg/l and 10^−4^ mg/l) and the mix of Fe_3_O_4_+SiO_2_ at the corresponding concentrations of 10^−3^ mg/l and 10^−4^ mg/l on the length of roots and seedlings, and the increase in the viability of plant cells under the influence of a stress factor (based on Evans blue staining). Field experience has shown the ambiguous effect of nano-printing of seeds on plant productivity.

## INTRODUCTION

Rapid and uniform germination is the key to agricultural production and can be achieved by seed “priming” methods (Gerna et al., 2018).

“Priming” is a well-known treatment for improving the quality of seeds. The seeds are primed showing an increase in germination, which leads to a high level of resistance to biotic / abiotic stress, yield and yield. All these characteristics, which increase the competitiveness of products, are directly related to seed energy, a complex of agronomic traits controlled by a variety of genetic and environmental factors (Paparella et al., 2015).

Priming of seeds with nanomaterials is an effective and development of equipment to increase strength of seedlings and the growth rates applied for certain types of plants. The small size and large surface area of nanoparticles allow them to demonstrate the unique physical, chemical and biological characteristics used in agriculture due to their high potential to improve seed germination and growth, plant protection, pathogen detection. The reaction of plants to nanoparticles depends on the plant species, its growth stage, and the nature of nanomaterials (Maroufi et al., 2011).

The most widely used NPs in agriculture are biogenic nanocrystalline compounds (Fe, Mo, Zn, Cu, Co, Se), due to their active participation in various redox processes and their presence in many enzymes and complex proteins (Quoc et al., 2014). In turn, Si is one of the most common macronutrients that have a positive effect on plant growth and development, but little information is available about its use for seed pre-treatment (Janmohammadi et al., 2015; Hoe et al., 2018; Hussain et al., 2019). Nano-iron has a high degree of bioavailability, which indicates its alternative use in living systems. Iron forms part of the catalytic centres of many redox enzymes, and also contributes to the formation of chloroplast proteins, which stimulate the development of the root and shoot systems (Kovalenko et al., 2006; Carvalho et al., 2018).

In the context of the above, the purpose of our research was to study the effect of colloidal solutions of nanoparticles of nutrients on the germination rate, oxidant status, yield and quality of products on the test plant *Pisum sativum*.

## MATERIALS AND METHODS

### Materials

The SiO_2_ NPs with a size of 30.7±0.3 nm and a ζ-potential of 27±0.12 mV were acquired from the company “Plasmotherm” (Moscow, Russia, http://plasmotherm.ru). Nanoparticles of Fe_3_O_4_ (80–100 nm, z-potential of 20 ± 0.14 mV) were acquired from the company “Advanced Powder Technologies” (Tomsk, Russia, www.nanosized-powders.com) (Figure 1).

**Figure 1 -.**
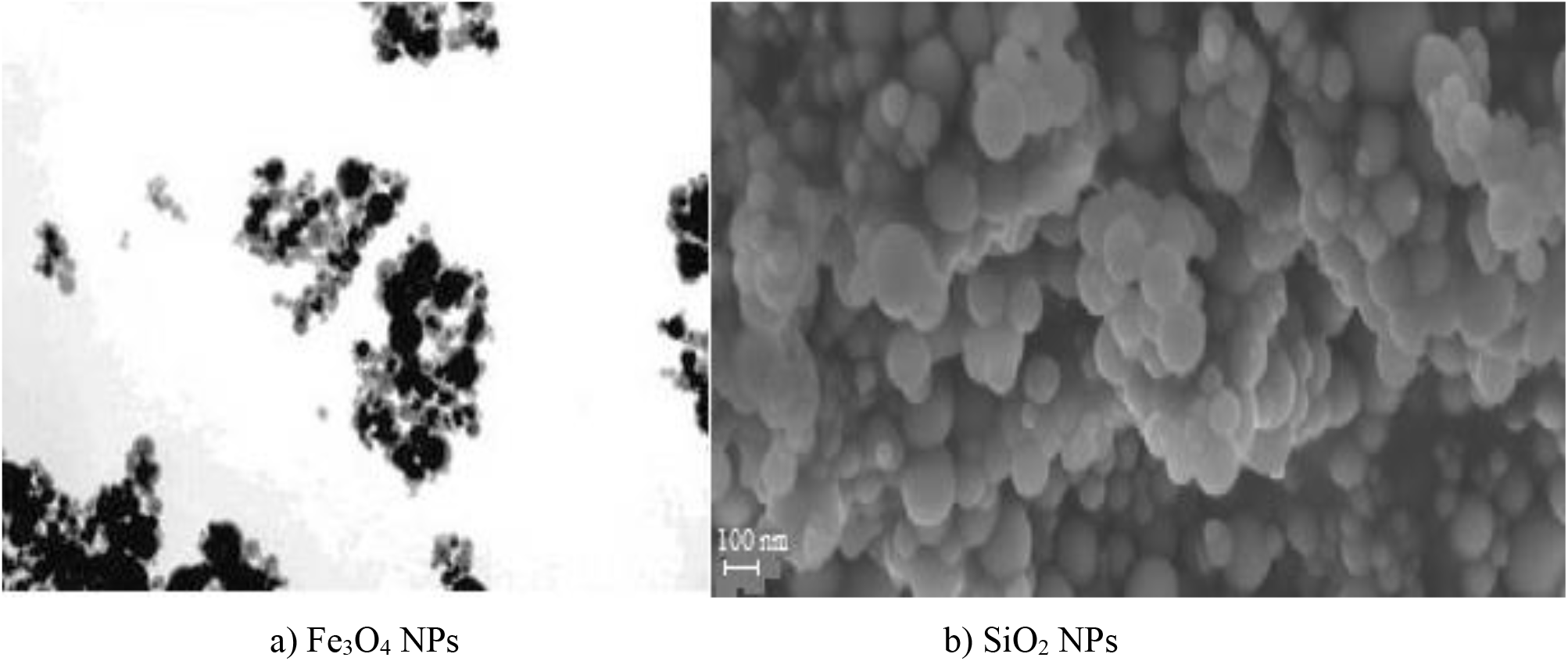
Electron microscopy of nanoparticles: a) transmission electron microscopy Fe_3_O_4_ NPs, b) scanning electron microscopy SiO_2_ NPs

For the preparation of NP solutions, exact amounts of the preparations were placed in glass flasks with distilled tap water and intensively dispersed by ultrasound at a frequency of 35 kHz for 30 minutes. Fold-dilutions were prepared to reduce the amount of nanomaterials. For seed treatment, the following concentrations were used: for Fe_3_O_4_ 10^−2^, 10^−3^ and 10^−4^ mg/l; for SiO_2_ 10^−2^, 10^−3^ and 10^−4^ mg/l; a mixed solution was obtained by mixing a suspension of Fe_3_O_4_ (10^−3^ mg/l) and SiO_2_ (10^−4^ mg/l) at a 1:1 ratio. The control version of the experiment included the treatment of seeds in deionized water.

To assess the efficiency of using nanoforms of iron and silicon, a traditional product for pre-sowing seed treatment was used (MivalAgro, acquired from AgroSil company (http://agrosil.ru)). The active substance of MivalAgro is a mixture of 760 g/kg of triethanolammonium salt of orthrocrekoxyacetic acid and 190 g/kg of 1-chloromethylsilatrane.

The pea variety «Flagship 12» was used as a test object. Seeds were provided by the FSSI «Samara RIoA» (Russia, http://samniish.ru), complied with all the requirements of the guidelines «The Order of biological assessment of effects of nanomaterials on plants using morphological characters» and were 1st class quality and not treated with disinfectant.

### Experimental design

The first phase of the experiment began with the pre-treatment of seeds of *P. sativum* in solutions of various agents (NPs SiO_2_, Fe_3_O_4_, MivalAgro) with subsequent drying in air. To determine the effective concentrations, the seeds were transferred to Petri dishes with 20 ml of deionized water.

Staining of plant root cells for loss of cell viability was carried out by staining freshly harvested roots from the control with Evans blue aquatic solution (0.25% v/v Evans blue, Sigma-Aldrich, Spain) for 10–15 min at room temperature followed by washing with CaCl_2_ solution (100 μm; pH 5.6) three times and visualization under a light microscope (Vijayaraghavareddy et al., 2017).

The second stage of the experiment involved the germination of plants in opaque plastic containers with soil in a climatic chamber. The containers measured 15 cm long, 12 cm wide and 10 cm high, and were filled with 2 kg of soil.

The soil was collected from an experimental field near the village Nezhinka Orenburg region of Russia (51°46’4”N55°22’7”E). The soil mass was collected from the 0–20 cm layer, dried, and the inclusions were selected and thoroughly mixed. Each container was sown with 20 pea seeds with germination in a climatic chamber (Pol-eko-1200 KK TOP+; Pol-EKO-Aparatura, Poland) at a relative humidity of 30%, air temperature of 25±2°C, and substrate temperature of 23±2°C. In total, the experiment involved 30 containers. Plant samples were collected and analysed 15, 25 and 35 days after emergence. The scheme of the experiment is shown in figures 2 and 3.

**Figure 2 –.**
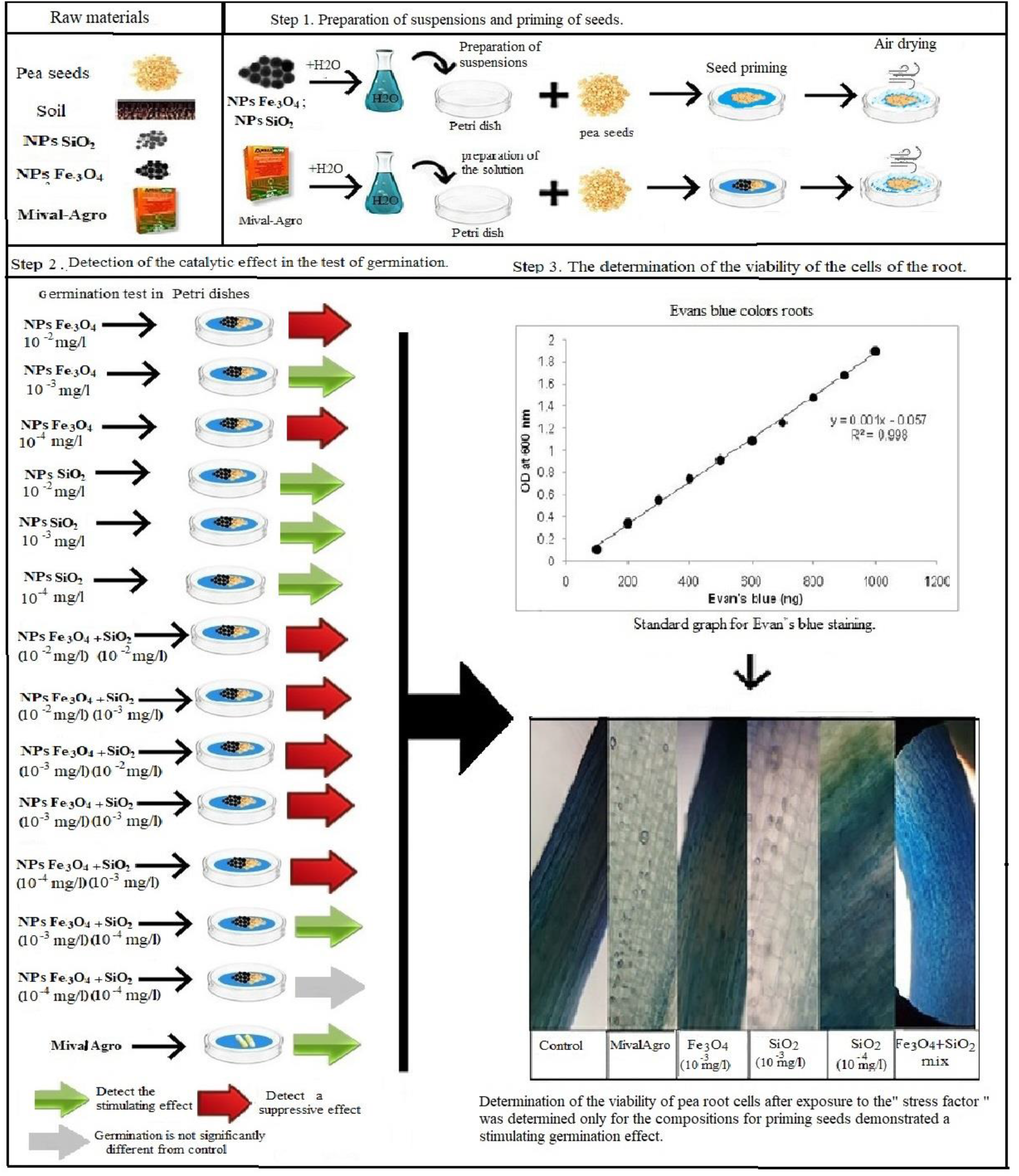
Experiment design (step 1, 2, 3)

**Figure 3 –.**
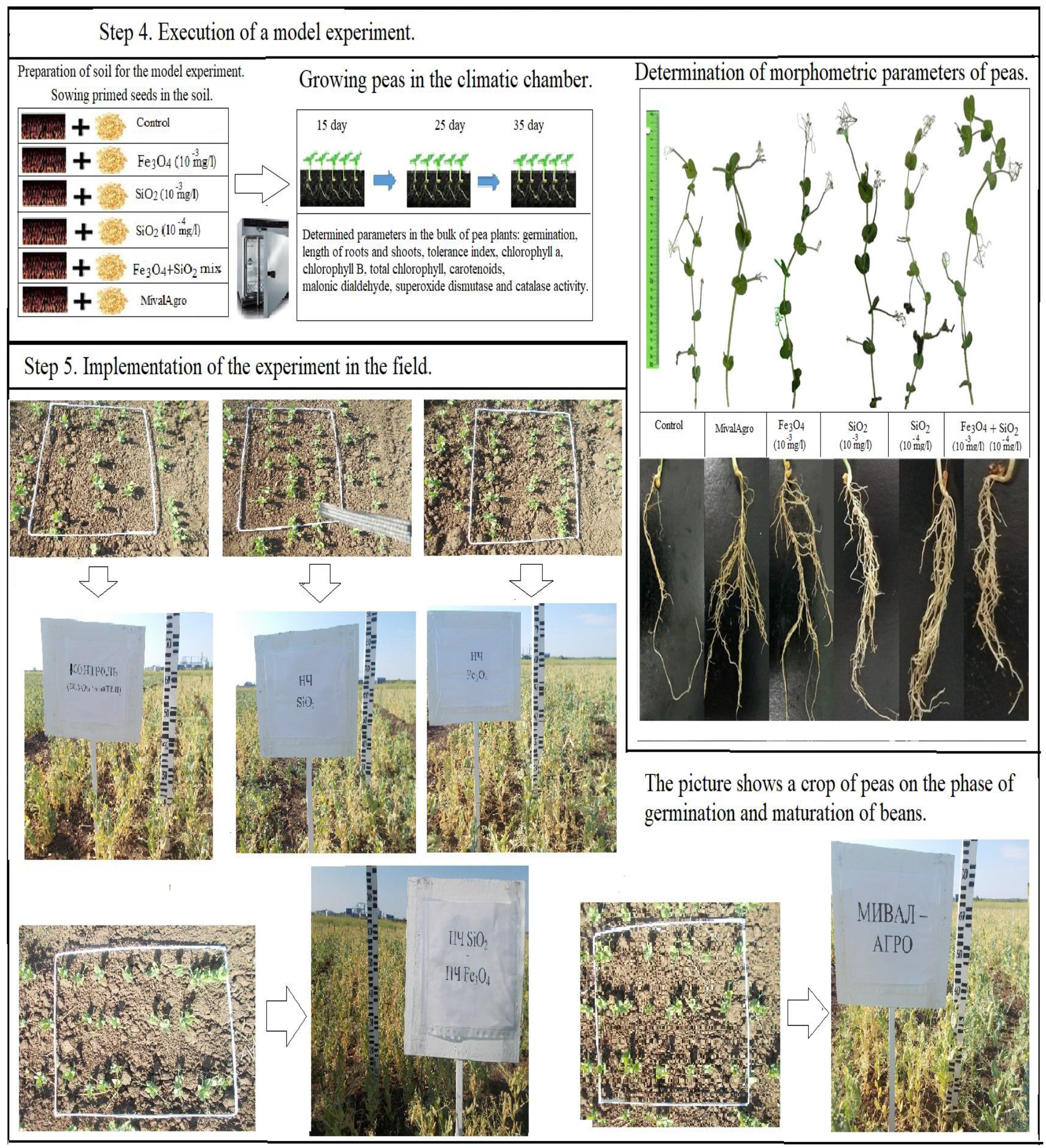
Experiment design (step 4,5)

### Field experiment

The experiments were conducted at the experimental site at village of Nezhinka (Orenburg region, Russia). The total size of the plot is 990 m^2^, which was further divided into 20 plots of 49.5 m^2^ (each measuring 1.65×30 m). The soil at the site is a Calcic Pachic Chernozem (IUSS Working Group WRB 2014). The soil characteristics are presented in Table 1.

Before sowing a two-step cultivation was carried out in order to destroy weeds, loosen and level the soil, and create a dense seed bed. The creation of favourable conditions for seed germination was achieved by rolling the field after sowing.

Pea seeds treated with a working solution of 10 l of agent per 1 ton of seeds were sown in the seed bed to a depth of 8 cm. As the growing conditions were very dry, and the contamination of the plots by weeds was minimal, no further care of the plots was necessary. The peas were harvested by combining (TERRION-SAMPO SR 2010) 82 days after sowing.

### Evaluation of vital, morphometric and biochemical parameters of test plants

There was a fixed germination percentage. The root and shoot lengths of seedlings were recorded using the standard centimetre scale. Measurements were carried out on roots after washing off the soil and drying on filter paper. The tolerance index (TI) of the plants was calculated (Wilkins, 1978).

The total activity of superoxide dismutase (SOD) was determined according to Giannopolitis and Rice (1972) (Bradford, 1976; Sibgatullina et al., 2011). Determination of catalase (CAT) was carried out using the method of Maehly and Chance (1955). The amount of lipid peroxidation products (LPO) in the soluble fraction of the homogenate and in the tissues was determined by the content of malondialdehyde (MDA), as described previously (Korotkova et al., 2017). Protein fragmentation was performed according to Chen and Bushuk (1970) and Bradford (1976).

Statistical analysis was performed using standard ANOVA techniques followed by the Tukey test (SPSS ver. 17.0). The Spearman method was used to determine the coefficient of correlation. Differences were considered statistically significant at p<0.05.

## RESULTS AND DISCUSSION

### Laboratory experiment

The laboratory experiment on the germination of seeds treated with 10^−3^ mg/l Fe_3_O_4_, 10^−3^ and 10^−4^ mg/l SiO_2_, and the mixture at a ratio of 1:1 (Figure 3) demonstrated reliable stimulation of germination with respect to the control by 10^−3^ mg/l and the mixture of Fe_3_O_4_ and SiO_2_, and demonstrated their effectiveness in relation to MivalAgro (Figure 4).

**Figure 4 –.**
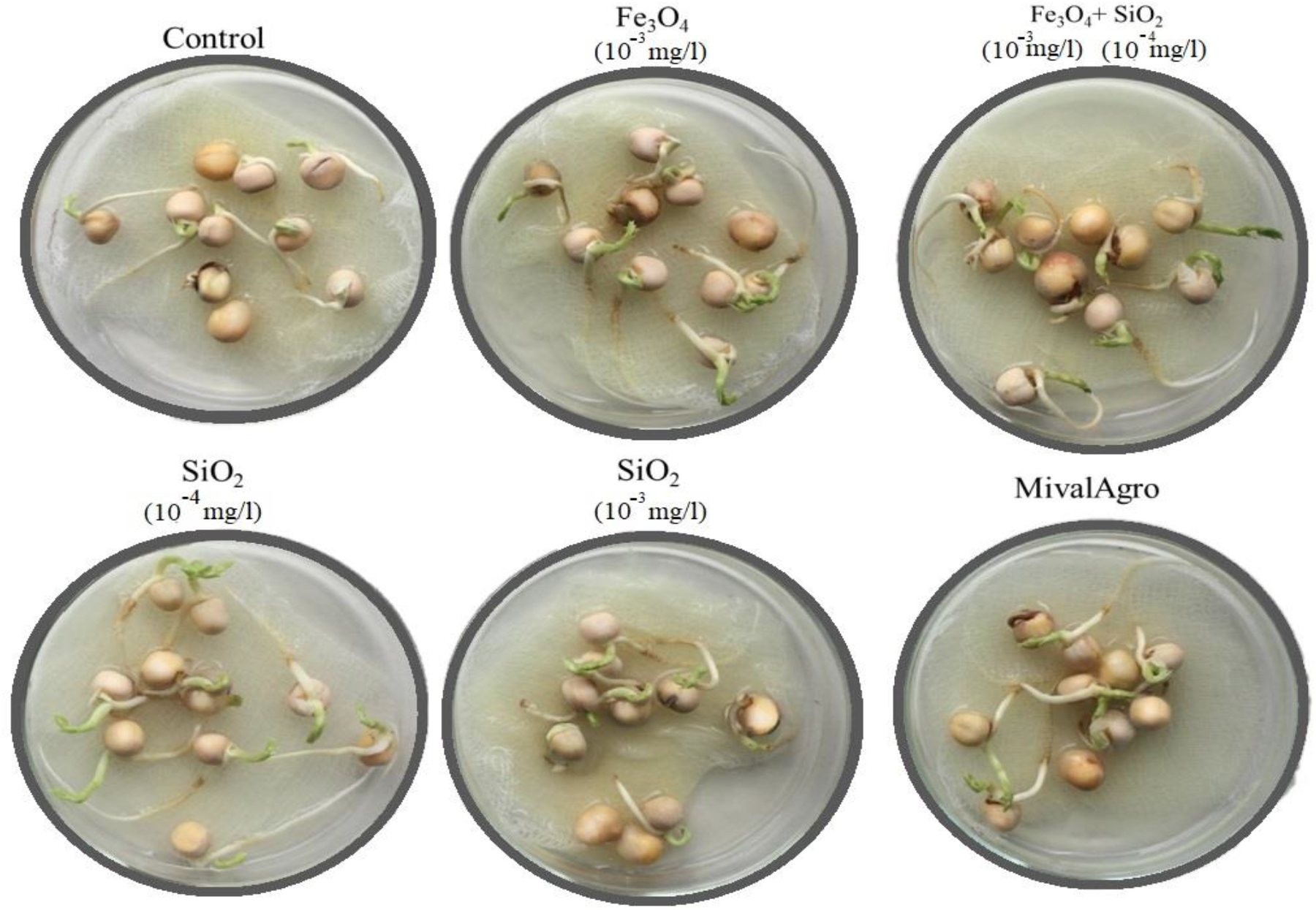
Pea seeds in germination dough

The study of the viability (V) of plant root cells was carried out using Evans blue dye, which is not able to penetrate into living cells but stains dead cells. The viability of pea root cells after exposure to the “stress factor” was determined only for those treatments having a stimulating effect on germination. Figure 5a shows a directly proportional increase in V after treatment of plants with 10^−3^ mg/l of the NPs SiO_2_ (up to 56.8 and 63.4% relative to plants in the positive control, 100% dead cells) and Fe_3_O_4_+SiO_2_ (up to 55%) (p<0.05).

**Figure 5 –.**
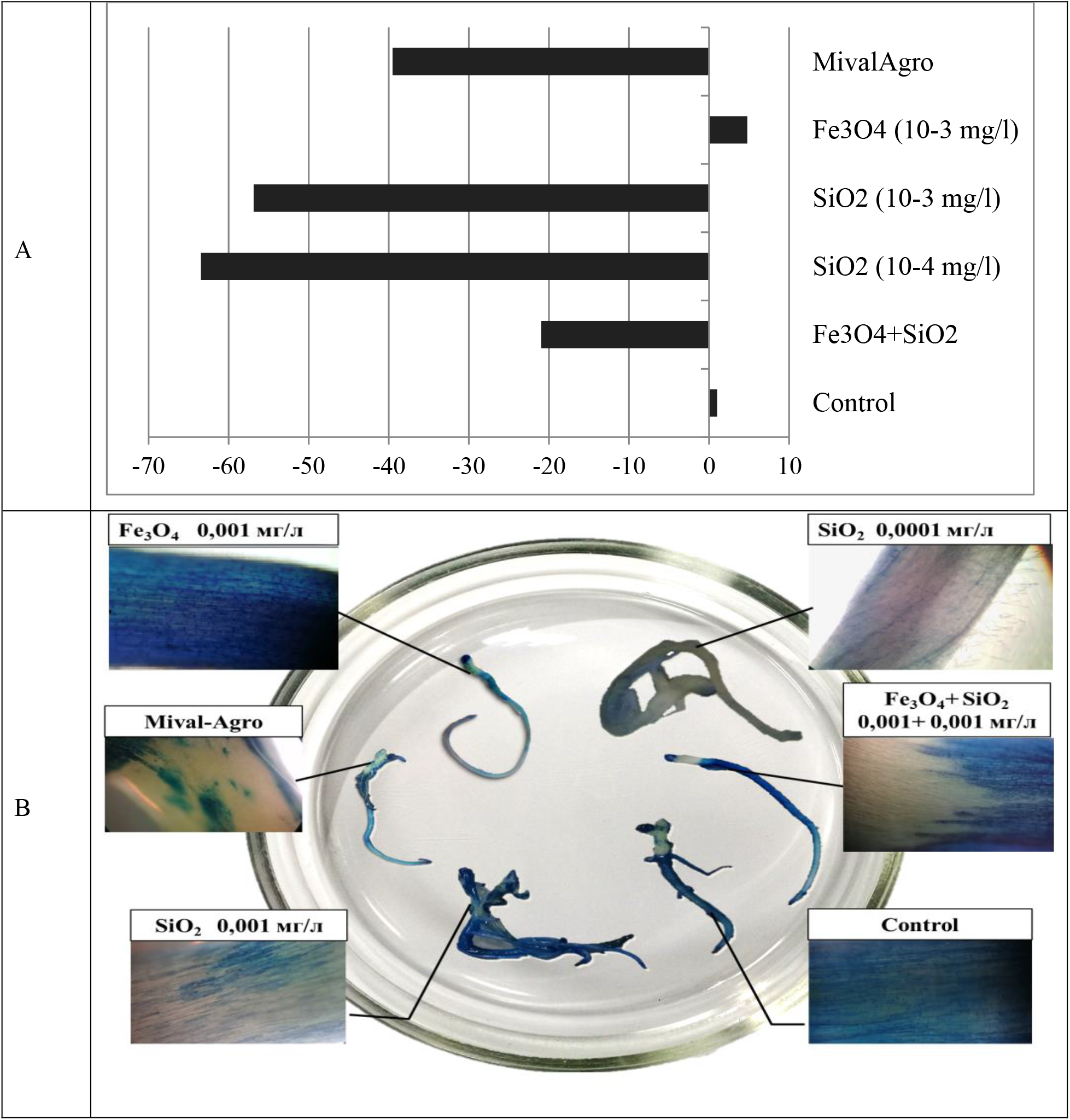
Viability of *P. sativum* roots after treatment of seeds with solutions of agents: A) diagram, % of control; C) percentage of living cells unstained with Evans dye after.… days of treatment: * variant significantly different from the control (value P≤0.05); C) cells with a damaged membrane in the elongation zone.

At the same time, the most pronounced protective effect against the impact of the stress factor on the root systems of plants was shown by SiO_2_ NPs at 10^−4^ mg/l. The data obtained were confirmed by microscopic assessment of the root elongation zone 7–14 days after germination (Figure 5B).

### Climatic chamber experiment

The experiment examining the germination of plants in soil under climatic chamber conditions showed that the maximum germination relative to the control sample was achieved in seeds treated with Fe_3_O_4_ NPs at 10^−3^ mg/l and Fe_3_O_4_+SiO_2_ at 10^−3^ mg/l and 10^−4^ mg/l (Figure 6). Note that the lowest level of germination was observed in seeds treated with MivalAgro (total ~78% relative to control).

**Figure 6 –.**
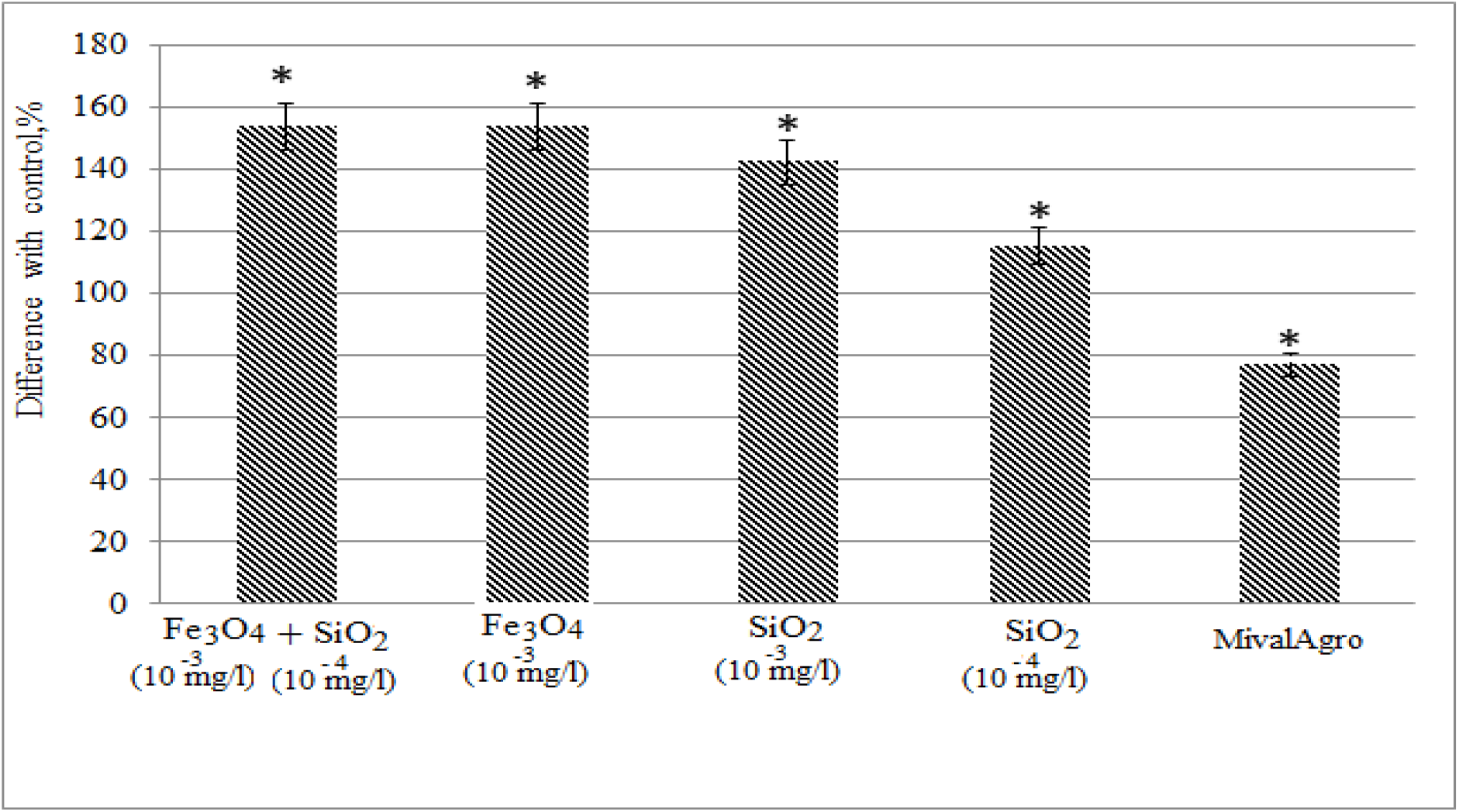
*P. sativum* germination. Data points with some / no symbols (*) represent no statistical significance at p≤0.05.

The stimulating effect of the NPs on *P. sativum* was most clearly manifested by the increased growth rate after germination. Thus, the studied compounds had a specific effect on the length of the seedlings, which significantly exceeded the control in the SiO_2_ treatment at 10^−3^ and 10^−4^ mg/l at all stages of the study (from 15 to 35 days), and after 25 days in the 10^−3^ mg/l Fe_3_O_4_ and Fe_3_O_4_+SiO_2_ treatment (Figure 7A).

**Figure 7 –.**
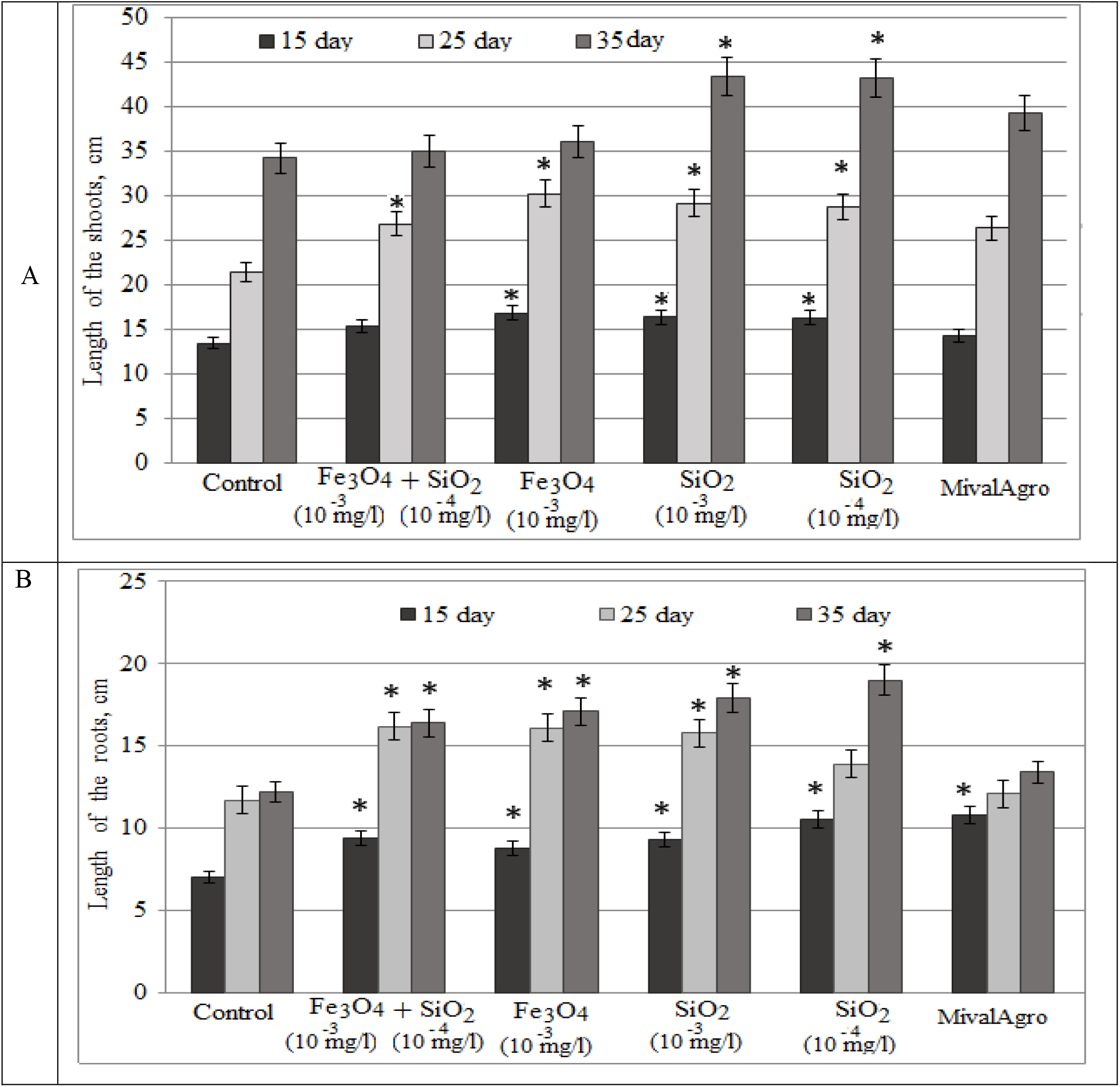
Growth parameters of *P. sativum* plants after exposure to various compounds: A - leaf length, B - root length. Bars are mean ± SEM (standard error of the mean). Data points with some/no symbols (*) represent no statistical significance at p ≤ 0.05.

MivalAgro significantly stimulated root growth only after 15 days of the experiment, and this stimulation was gradually offset in subsequent periods of growth. At the same time, it had no significant effect on the length of the above-ground organs of peas. Similar results were obtained regarding the length of pea roots, with maximum development in treatments with SiO_2_ solutions at different concentrations, 10 mg/l Fe_3_O_4_ and Fe_3_O_4_+SiO_2_ (Figure 7B).

The integral metric characteristic of model plants in response to exogenous stimulators (TI) is presented in Figure 8. A significant effect of seed treatment on the TI of plants at all stages of the study was observed at both concentrations of SiO_2_, and after 15–25 days for Fe_3_O_4_ combined with SiO_2_. The obtained data are consistent with the literature (Nazaralian et al., 2017; Yuan et al., 2018).

**Figure 8 –.**
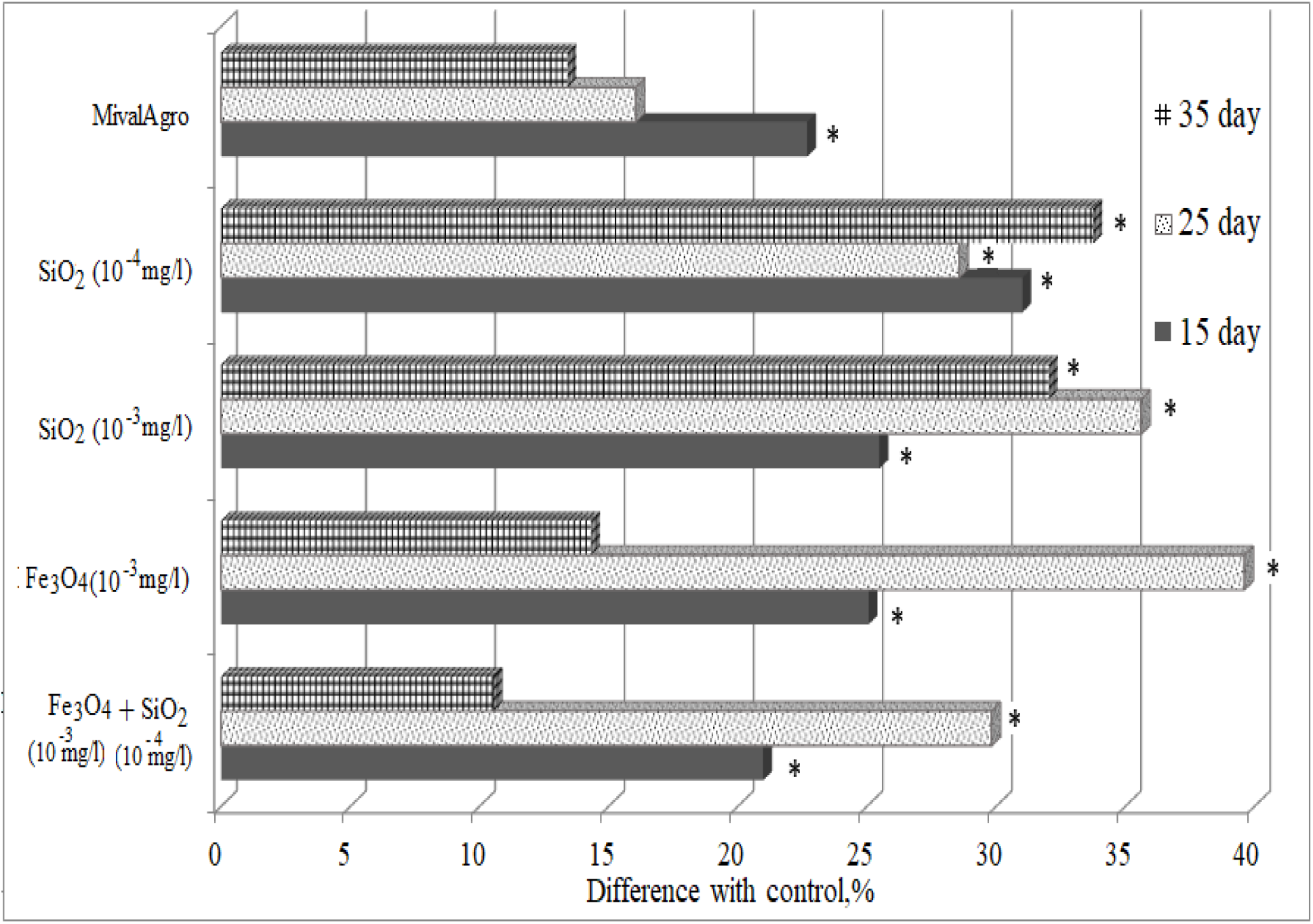
Tolerance index of *P. sativum* plants after exposure to various compounds. Bars are mean ± SEM (standard error of the mean). Data points with some/no symbols (*) represent no statistical significance at p ≤ 0.05.

The beneficial effects of SiO_2_ reported by Tahir and his colleagues (2010) were associated with its hydrophilicity (Romero-Aranda et al., 2006), and in experiments with wheat led to a significant increase in biomass and yield.

The results of the experiment showed that the enzymes SOD and CAT in peas react differently to the presence of the studied compounds (Figure 9). Thus, catalase activity in seedlings was 50–60% lower than that of the control in the first two weeks and on the 25th day there was an increase in CAT to the level of control plants. In turn, a sharp increase in the index was noted on the 35th day of exposure to the treatment agents, with the exception of MivalAgro, which indicates the exhaustion of the existing enzyme pool. Thus, an increase in CAT was recorded after SiO_2_ treatment at both concentrations (by up to 83% and 146%), Fe_3_O_4_ (by up to 111%) and Fe_3_O_4_+SiO_2_ (by up to 47%) (p<0.05).

**Figure 9 –.**
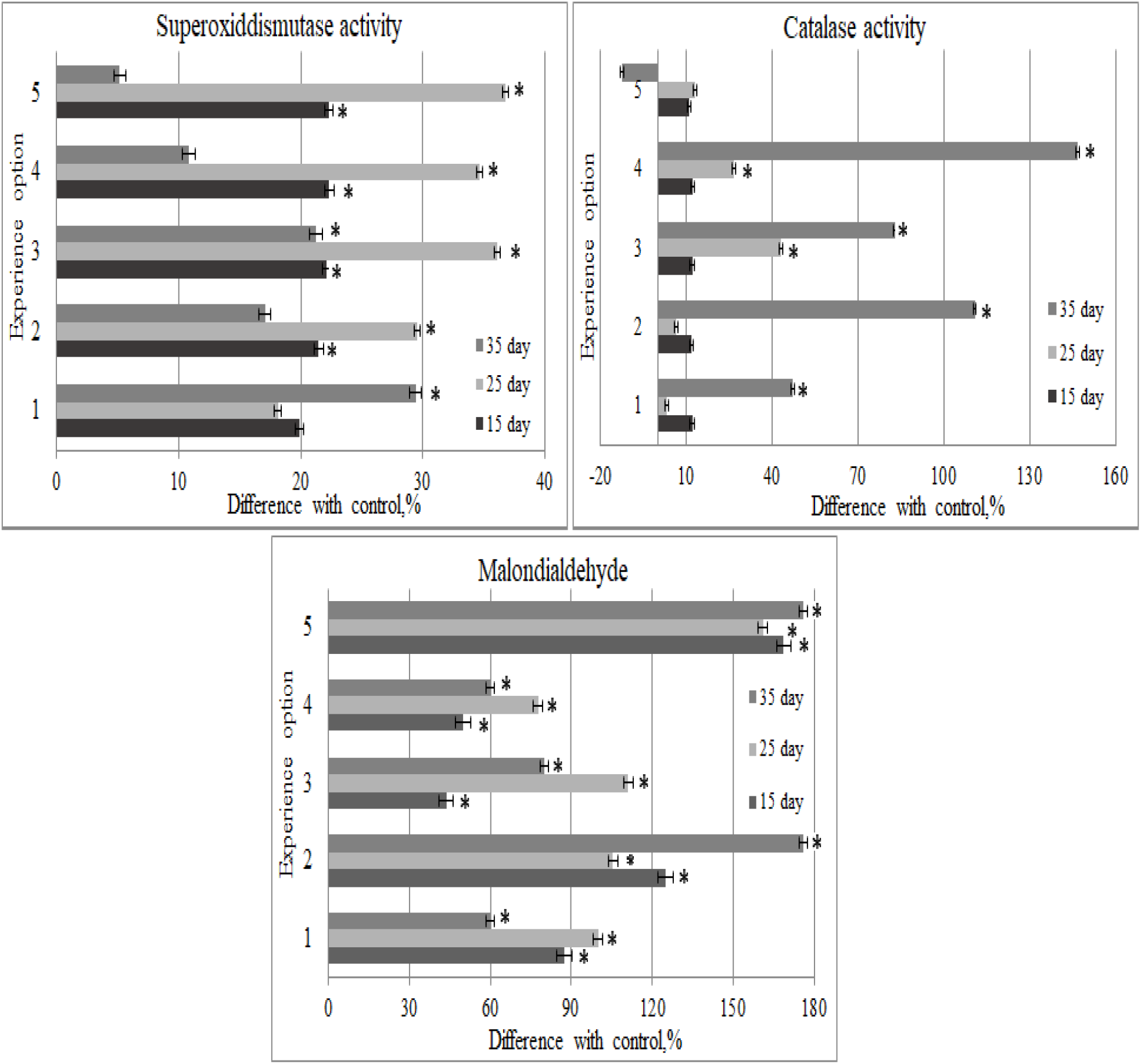
The activity of antioxidant enzymes (CAT, SOD) and the degree of lipid peroxidation (according to the MDA level) in *P. sativum* after exposure to various compounds: 1-Fe_3_O_4_+SiO_2_; 2 - Fe_3_O_4_ 10^−3^ mg/l; 3 -SiO_2_ 10^−3^ mg/l; 4 - SiO_2_ 10^−4^ mg/l; 5 – MivalAgro. Bars are mean ± SEM (standard error of the mean). Data points with some/no symbols (*) represent no statistical significance at p ≤ 0.05.

CAT is the main antioxidant in plants, preventing their oxidative damage and acting as a neutralizer of radicals. Our results show that SiO_2_ NPs can cause oxidative stress only at concentrations above 200 mg/l. Similar results have been published previously (Yang et al., 2012; Wang, Zhang, 2014).

Differences in stress-dependent total SOD activity were less noticeable. A slight increase in enzyme levels (over 30%) relative to the control was recorded in plants after 25 days of exposure to MivalAgro and SiO_2_ (p<0.05). At the same time, on the 35th day, the level of SOD decreased to the control value, which indicates the effective neutralization of superoxide radicals and intensification of CAT activity.

As a biological indicator of the development of oxidative stress in plants, the product of lipid peroxidation (LPO) of cell membranes – malondialdehyde (MDA) – was used. Free radicals are able to initiate LPO, as a result of which the membranes become permeable to ions and organic acids. Analysis of the degree of LPO in the seeds showed that pretreatment with nanoparticles had a multidirectional effect depending on the phase of growth. Thus, on the 25th day of the experiment, there was a small increase in the MDA level compared to the control in the case of treatment with 10^−3^ and 10^−4^ mg/l SiO_2_ and a mixture of Fe_3_O_4_+SiO_2_. However, this indicator was much lower than the level recorded after treatment with MivalAgro (1.5–2 times lower). Under experimental conditions, even within 35 days, the less strong accumulation of MDA in SiO_2_ and Fe_3_O_4_+SiO_2_ treatments (not more than 40%) indicates the resistance of plants to the nanoform of these elements.

In contrast, plants treated with Fe_3_O_4_ had a slightly higher content of MDA from 15 to 35 days, comparable to the drug MivalAgro i.e. up to 176% (p<0.05). These results show that nanoparticles can also lead to the formation of free radicals and damage the integrity of the membrane.

It has previously been reported that SiO_2_ nanoparticles cause oxidative stress in plants and lead to LPO (Slomberg et al., 2012). Our results are consistent with other studies indicating an increase in LPO in different plant species after SiO_2_ treatment (Yang et al., 2012; Wang, Zhang, 2014).

Thus, the activity of SOD and CAT can be considered an important element of antioxidant protection of plant cells from stress factors, including excess or lack of trace elements. However, under experimental conditions, the activity of these enzymes with a small increase in MDA indicates the stress of a redox imbalance due to mineral deficiency caused by the lack of nutrients in the growing medium.

### Field experiment

The weather conditions prevailing during the growing season of the *P. sativum* were extremely unfavourable: extremely high air temperature (30–40 C) accompanied by a high daily deficit of air humidity from 16 to 22 MB. The negative impact on the grain yield was caused by the lack of productive soil moisture due to the low snow-cover during winter and deep freezing of the soil. During the growing season only 38 mm of precipitation fell, which is 39% of the mean annual values.

The moisture supply during the period from germination to full ripeness was only 28.4% of the level of demand. Due to the negative influence of the above factors, the grain yield was low (1.9–2.6 c/ha), while the difference between treatment groups was insignificant (Table 2).

A tendency for a decrease in productivity under the influence of the treatments was noted, except for the low-concentration SiO_2_ treatment, in which grain yield was higher than in the control (without treatment) by 0.1 c/ha.

Analysis of the components of grain yield showed that the number of beans on the plant, seeds in the bean, and grain weight were at or below the control level in the treatment groups. The positive impact of the treatment of SiO_2_, Fe_3_O_4_ and organic silicon (MivalAgro) manifested itself in the form of an increased density of 5–7 plants per 1 m^2^ compared to the control variant.

Analysis of the proteins in the plant after treatment with exogenous stimulators in the field showed a marked increase in the albumin pool after exposure to a mixture of Fe_3_O_4_+SiO_2_ 88% (Figure 10). The content of globulins decreased to 9.8% relative to the control.

**Figure 10 –.**
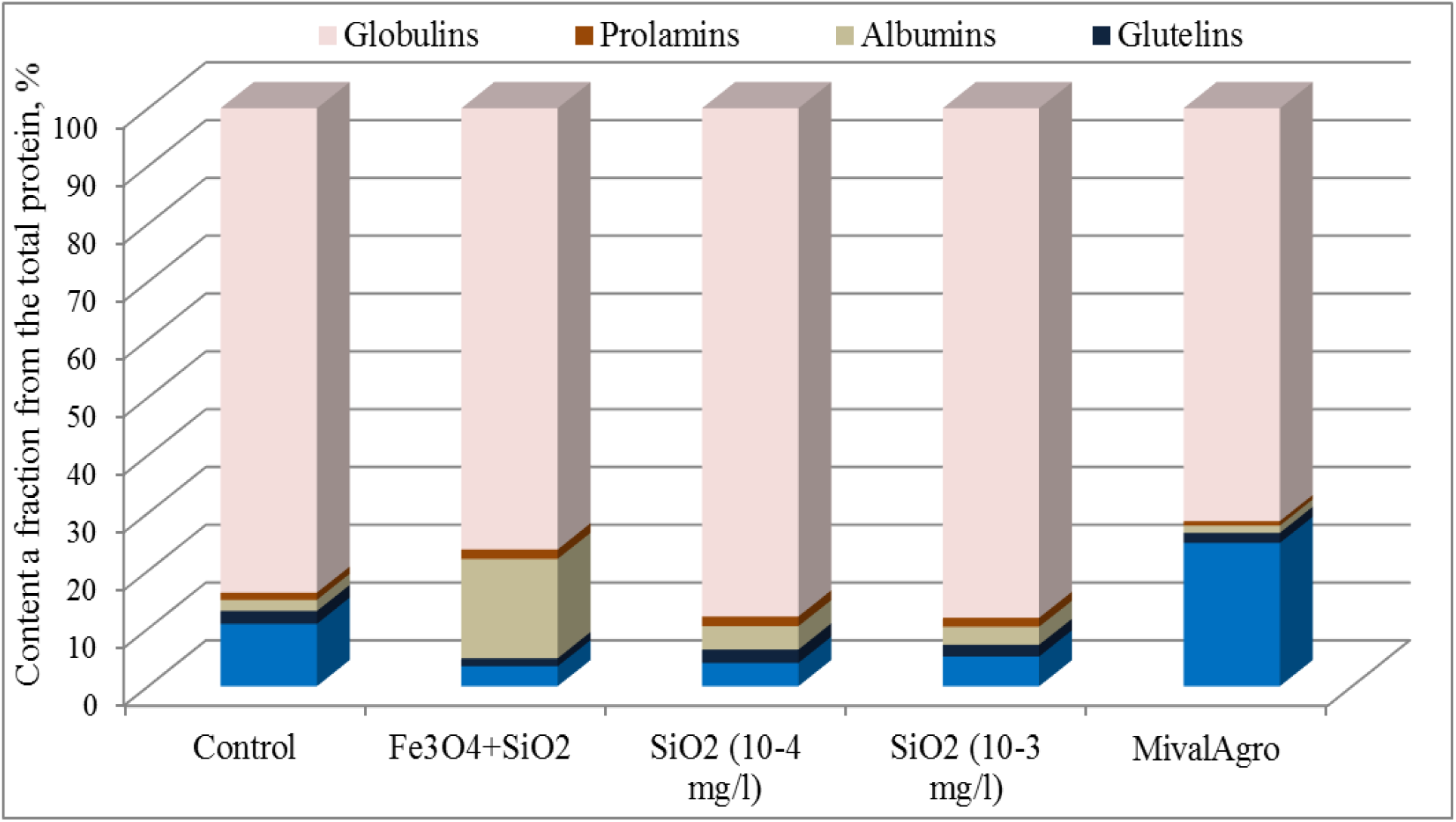
Protein content of *P. sativum* seeds. Data points with some/no symbols (*) represent no statistical significance at p ≤ 0.05.

Treatment with SiO_2_ led to a small increase in the amount of globulins relative to the control (no more than 5%) while reducing the fraction of gluten. It should be noted that there was a marked increase in the amount of gluten after exposure to MivalAgro (by 50%), which was probably caused by specific activation of protein metabolism.

A decrease in prolamine and glutelin in rice grown in medium with nanoparticles of CeO_2_ was previously reported (Rico et al., 2013). A study by Zhao and colleagues (2014) showed that ZnO NPs did not affect protein fractions in cucumber, but CeO_2_ at a dose of 400 mg/kg increased the amount of globulins and reduced the amount of gluten. Thus, the effect of treatment of seeds with nanoparticle suspensions on the composition of the protein complex has no element and/or species-specific character.

## CONCLUSION

The influence of colloidal solutions of nanoparticles of Fe_3_O_4_, SiO_2_, and their mixture at a ratio of 1:1 on the complex of physiological and biochemical parameters of the plant Pisum sativum was studied. The study was based on laboratory and field experiments, from which the following conclusions can be drawn:

1. The comparison of different compositions of nano compounds with the organic analogue MivalAgro in the laboratory showed that reliable stimulation of seed germination and growth of seedlings was achieved by the NPs Fe_3_O_4_ (10^−3^ mg/l), SiO_2_ (10^−3^ and 10^−4^ mg/l) and their mixture (Fe_3_O_4_+ SiO_2_).
2. After more than 25 days, SiO_2_ at 10^−4^ mg/l and Fe_3_O_4_ showed a stimulating effect on plant growth relative to MivalAgro.
3. The results of the field experiment did not demonstrate an increase in plant resistance to environmental stress factors, which indicates the need for further research on the introduction of nanotechnological solutions in crop production.

## Acknowledgment

The research was supported by the Ministry of Science and Higher Education in accordance with the state assignment for Ural State Mining University No. 0833-2020-0008 ‘Development and environmental and economic substantiation of the technology for reclamation of land disturbed by the mining and metallurgical complex based on reclamation materials and fertilizers of a new type’. We obtain the scientific results with the staff of Center for the collective use by using funds of the Center for the collective use of scientific equipment of the Federal Scientific Center of biological systems and agricultural technologies of RAS as well (No Ross RU.0001.21 PF59, the Unified Russian Register of Centers for Collective Use - http://www.ckp-rf.ru/ckp/77384).

